## Supplementary Figures and legends for "TopBP1 utilises a bipartite GINS binding mode to support genome replication": Supplementary information and legends.pdf

### Supplementary Figure 1

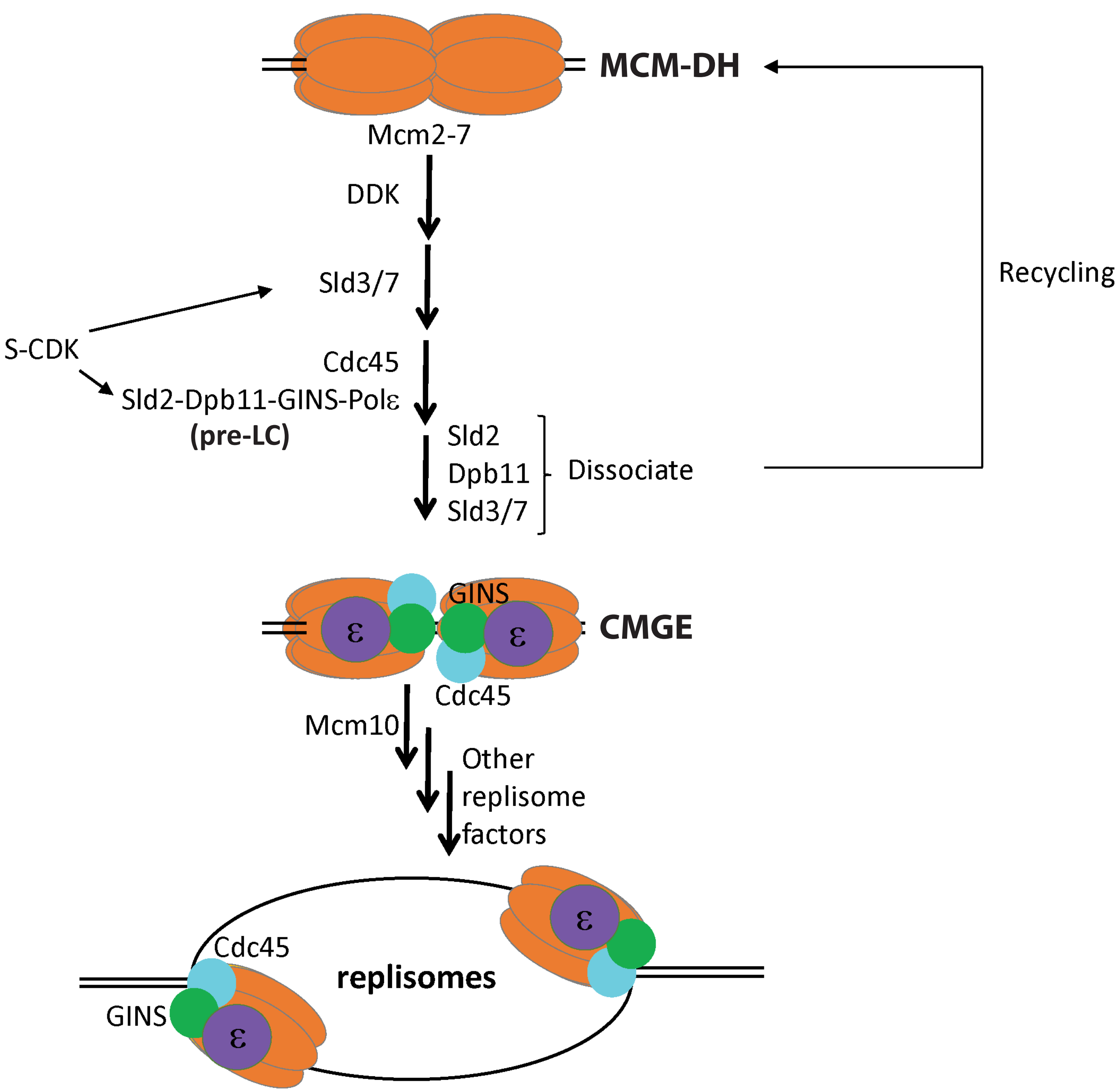

### Supplementary Figure 2

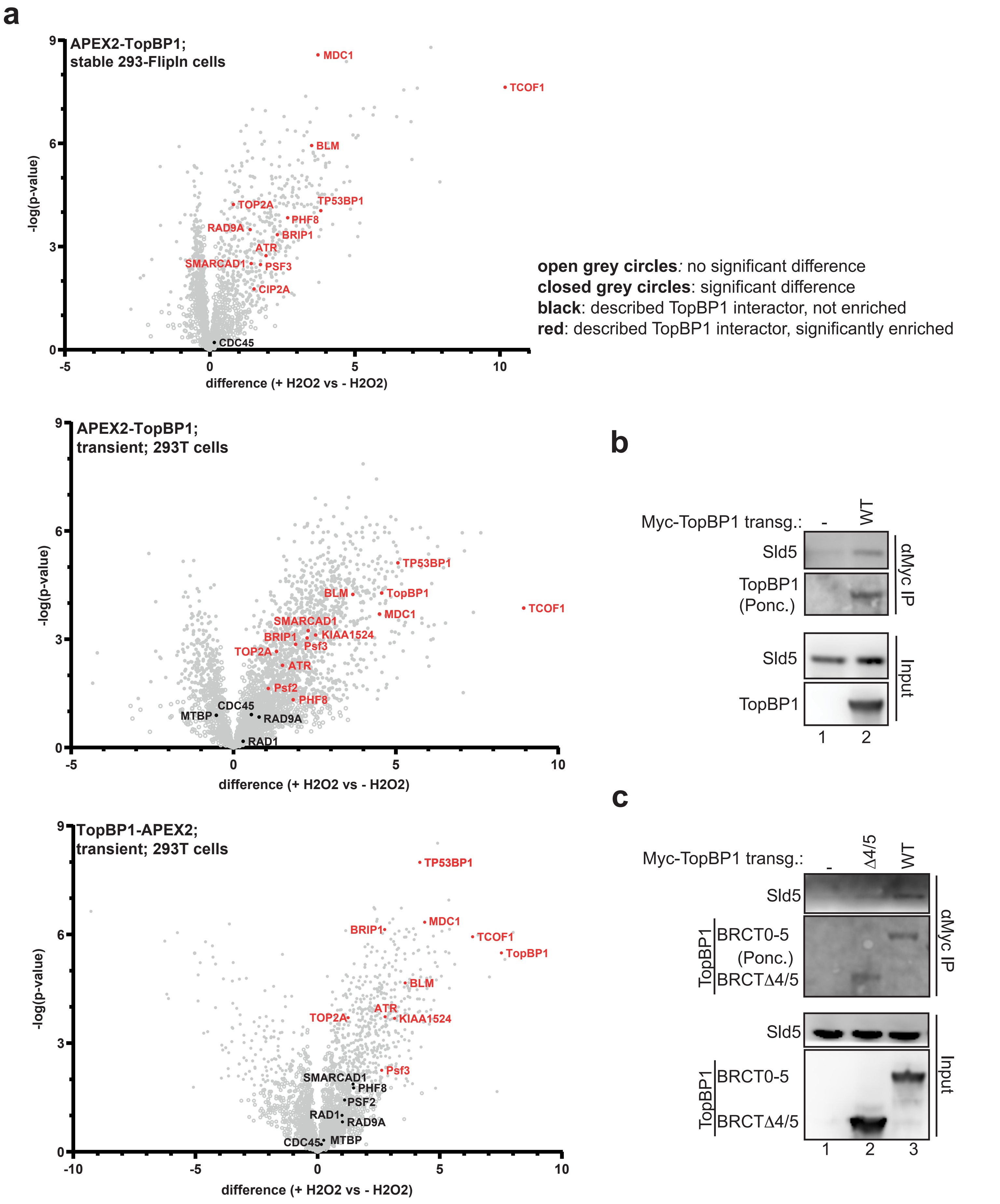

### Supplementary Figure 3

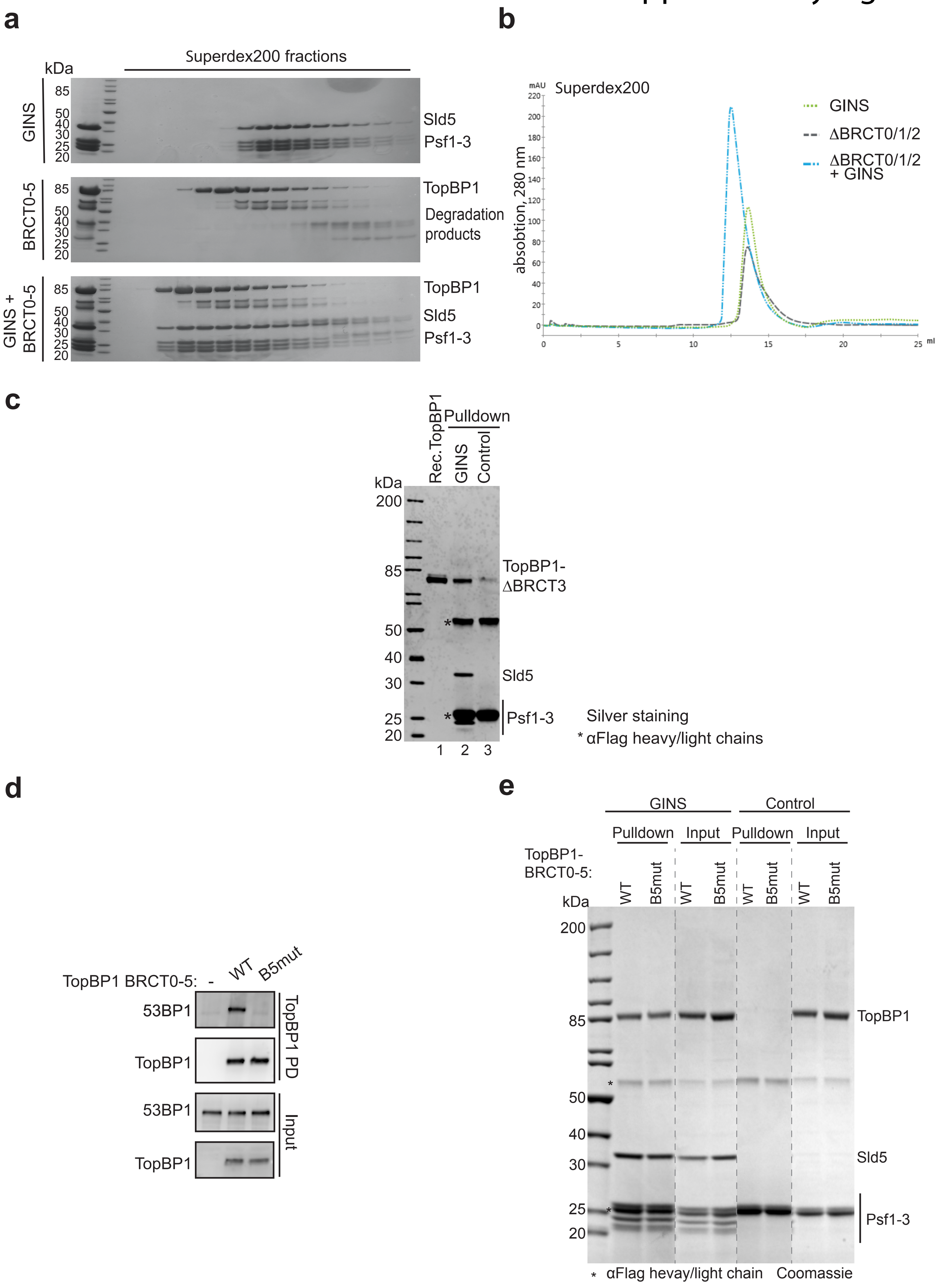

Supplementary Figure 4

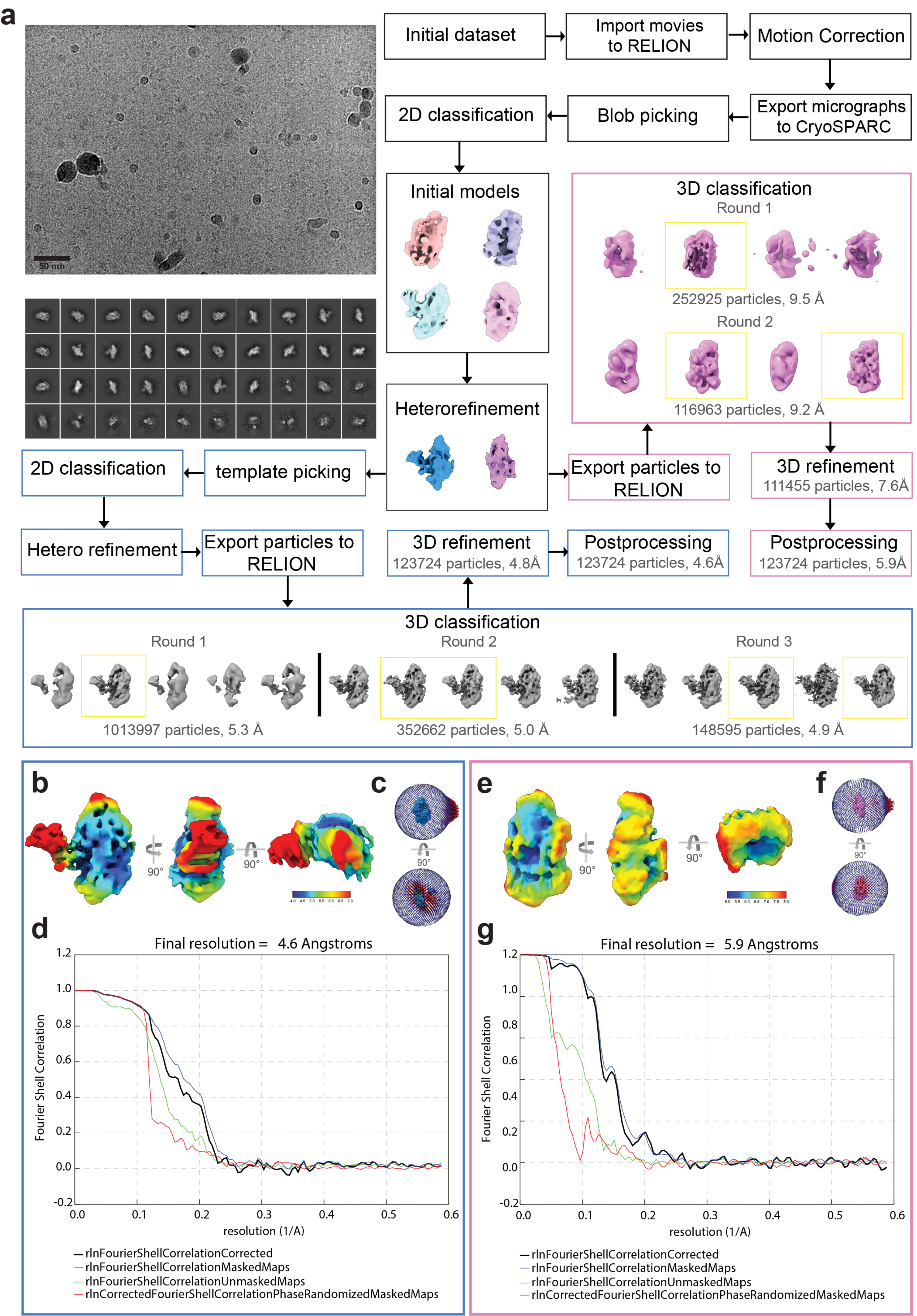

### Supplementary Figure 5

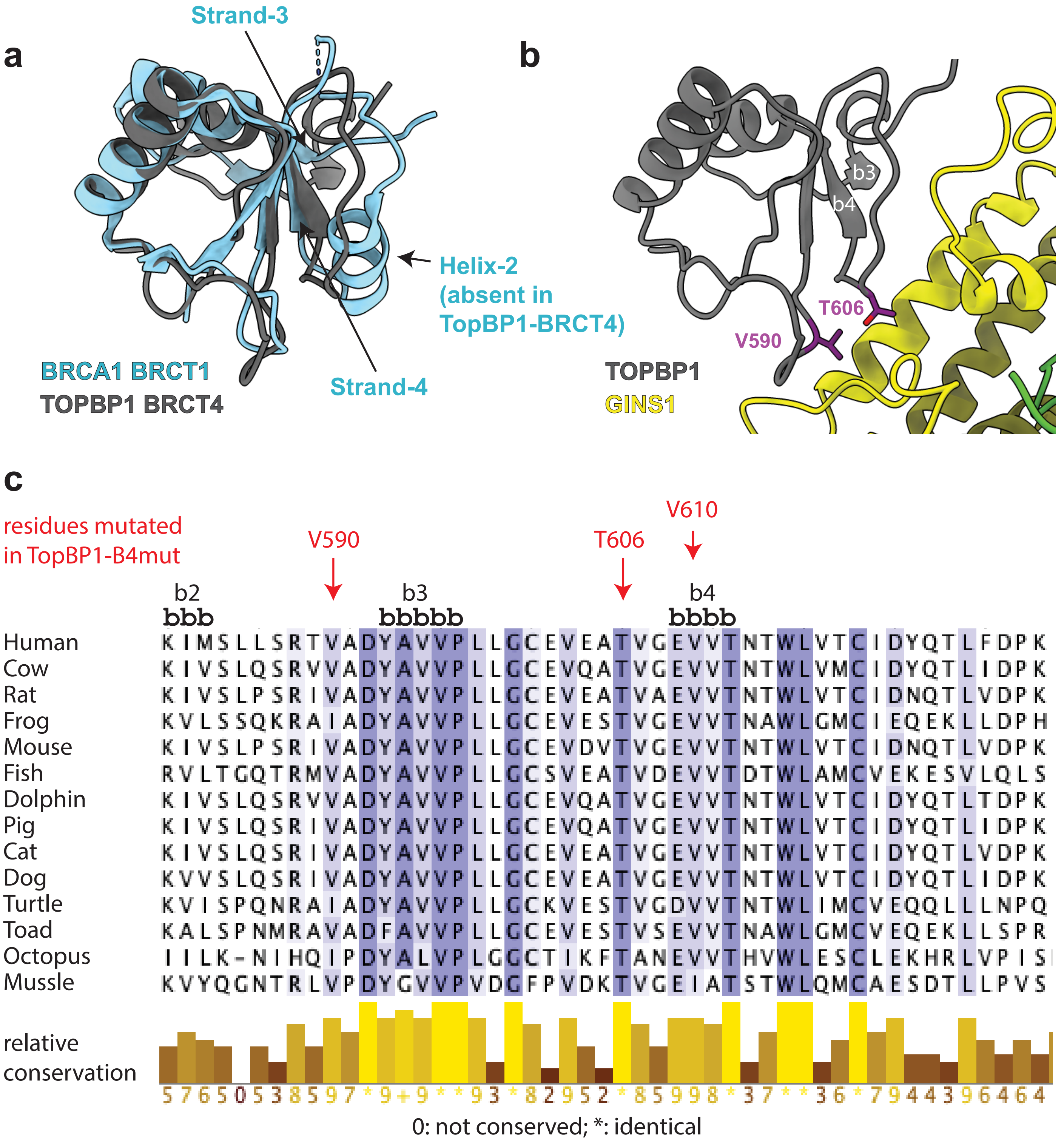

### Supplementary Figure 6

**a**

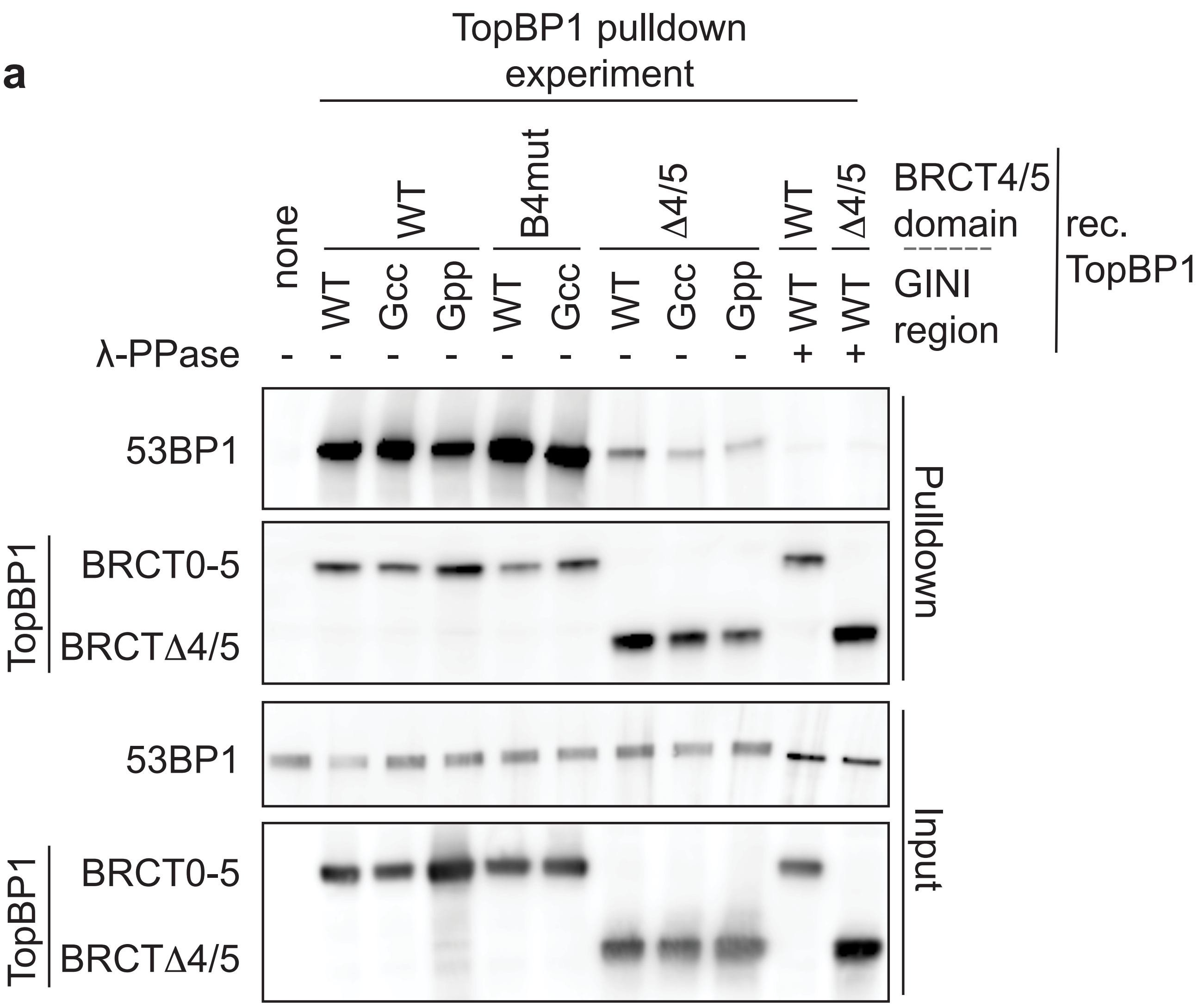

**b**

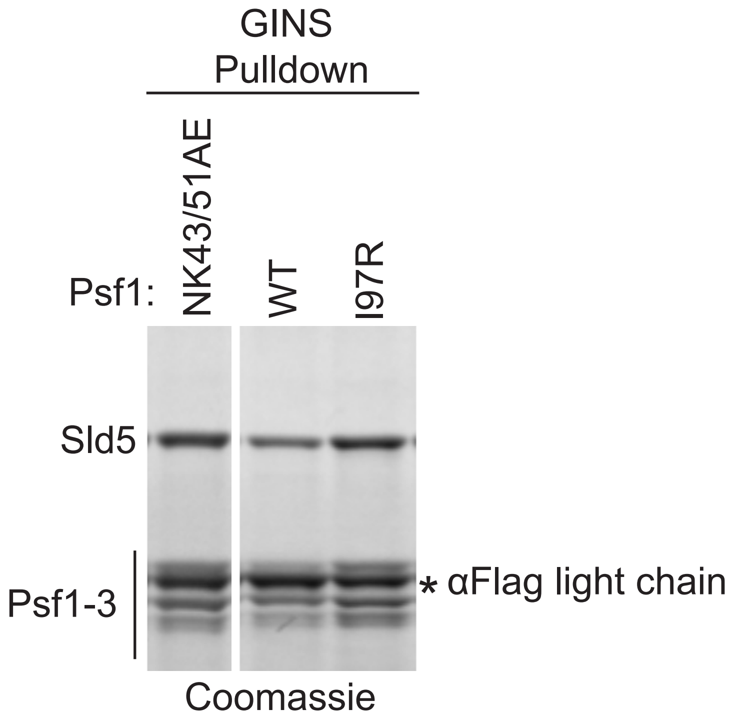

### Supplementary Figure 7

TopBP1 BRCTs 0-5

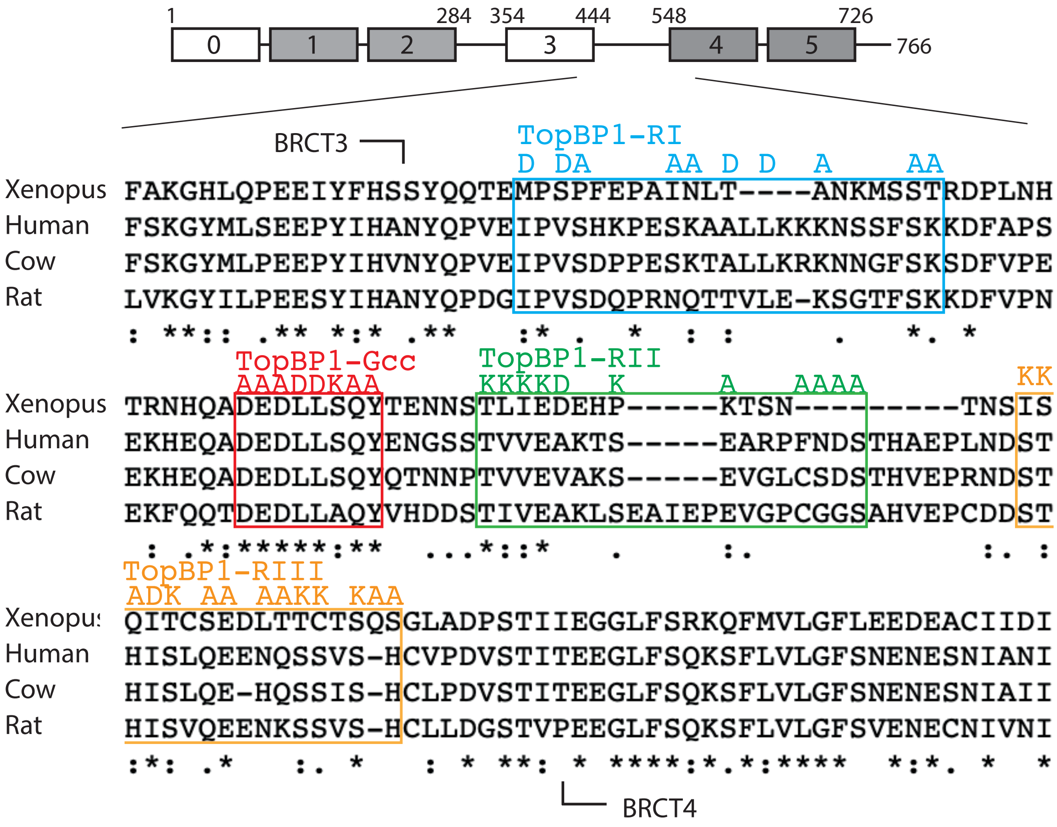

### Supplementary Figure 8

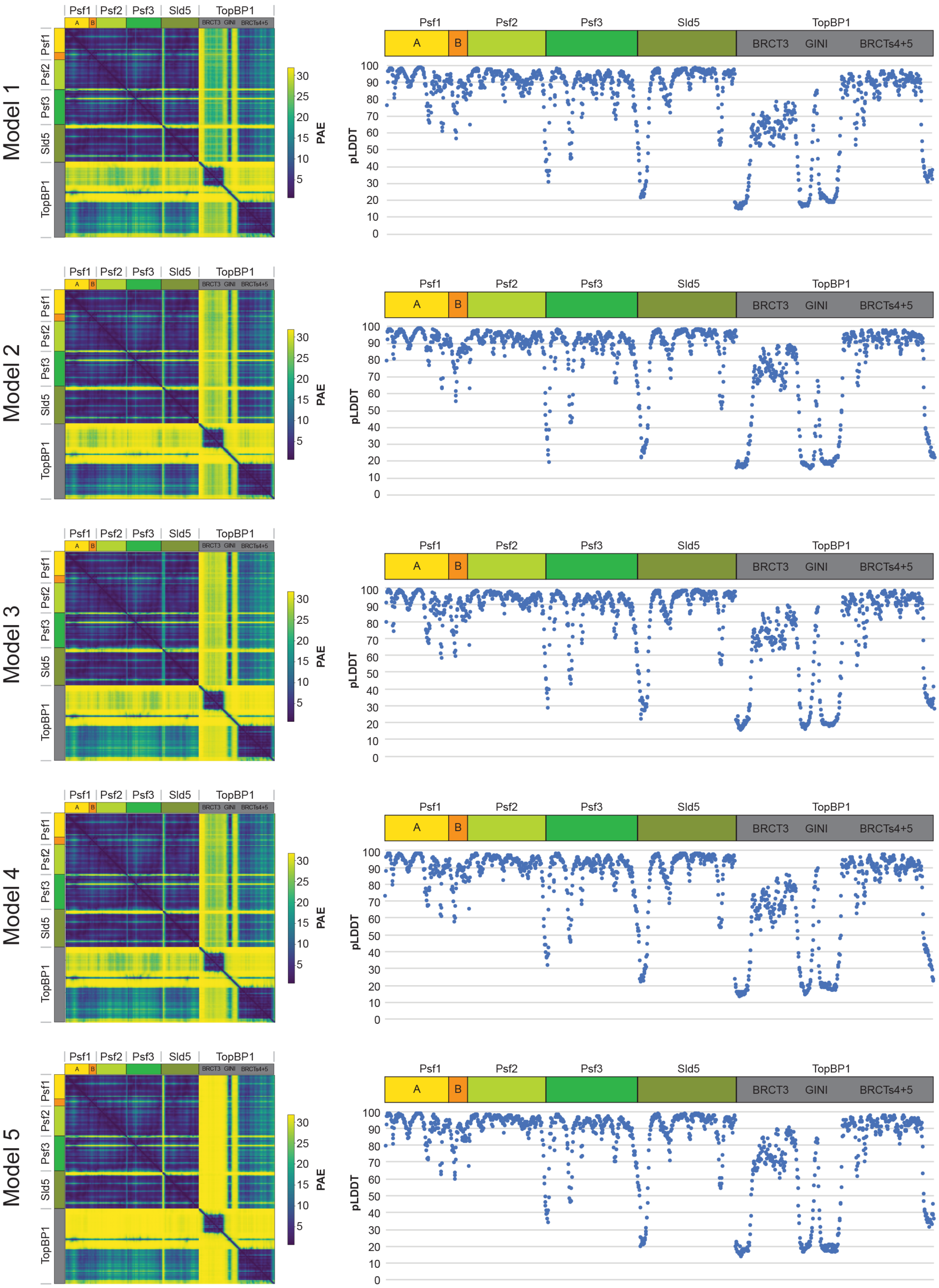

### Supplementary Figure 9

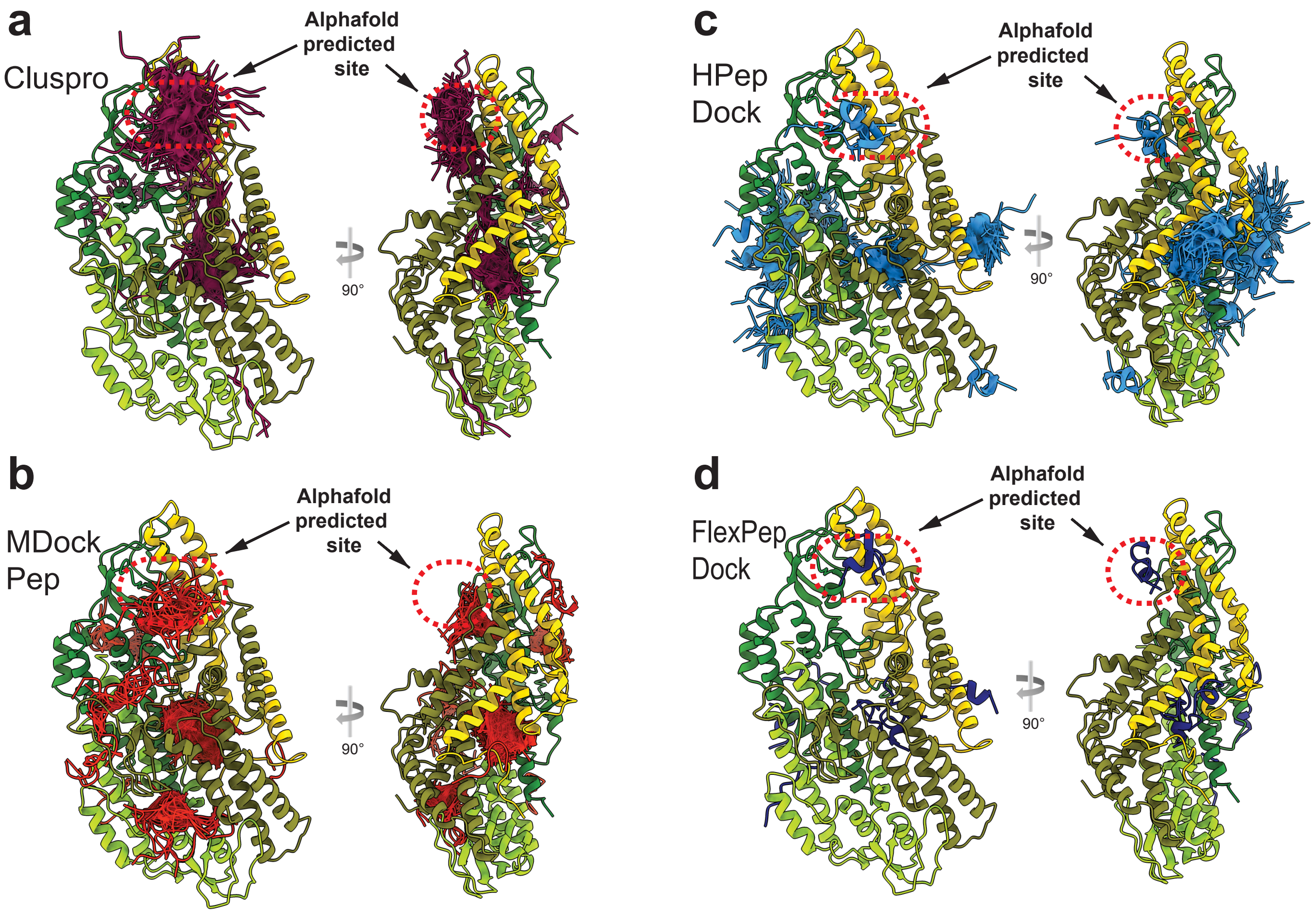

### Supplementary Figure 10

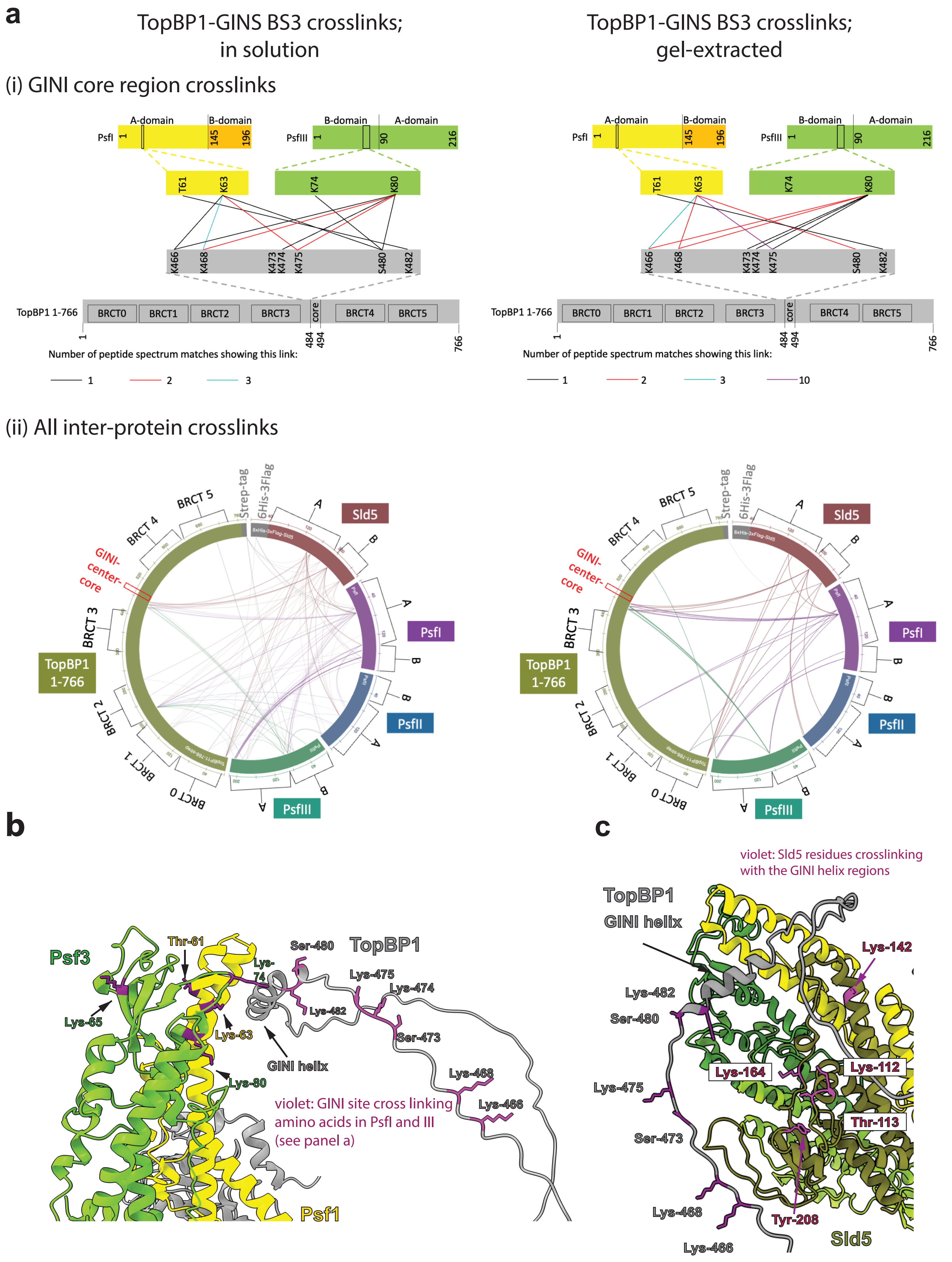

### Supplementary Figure 11

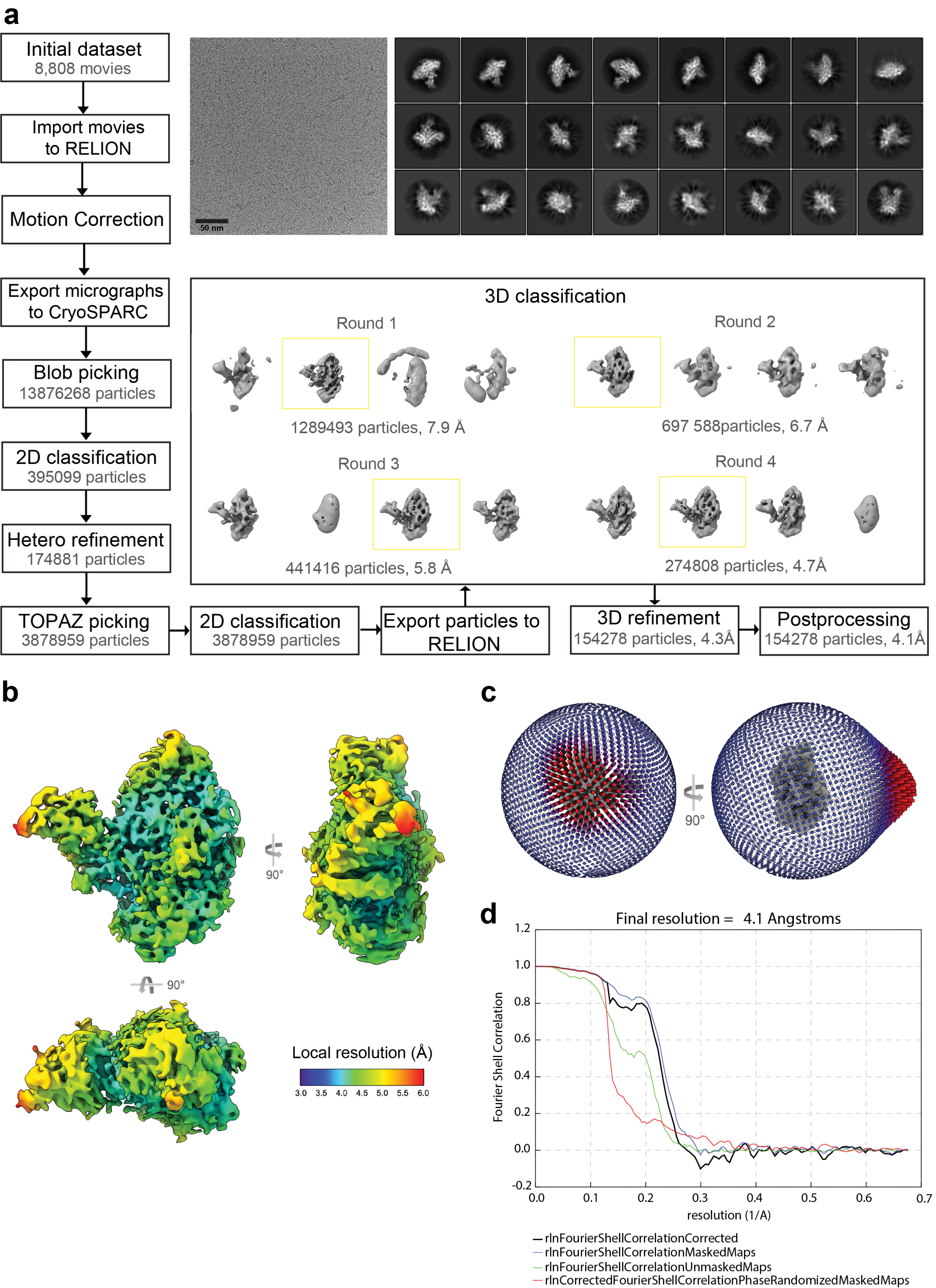

### Supplementary Figure 12

**a**

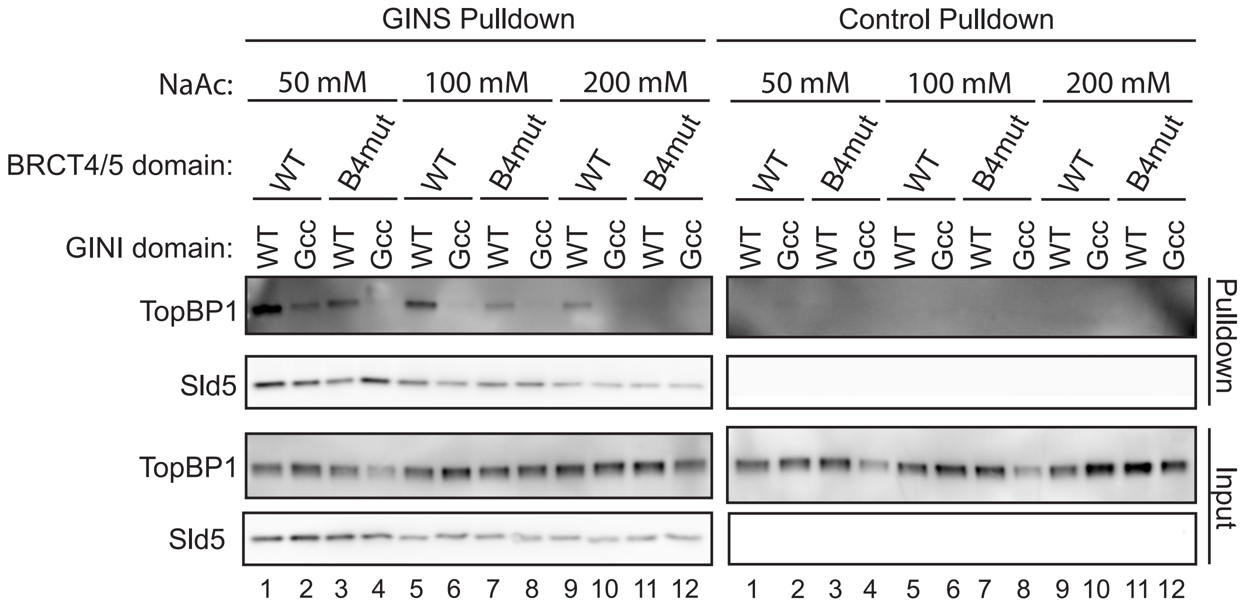

**b**

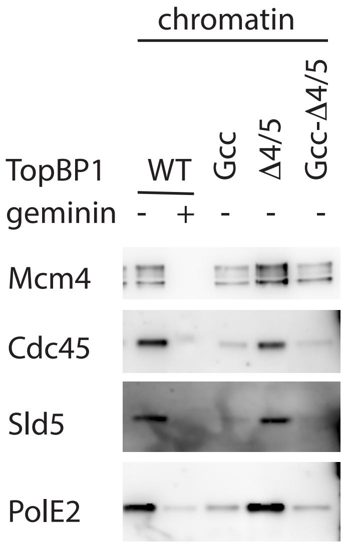

### Supplementary Figure 13

**a**

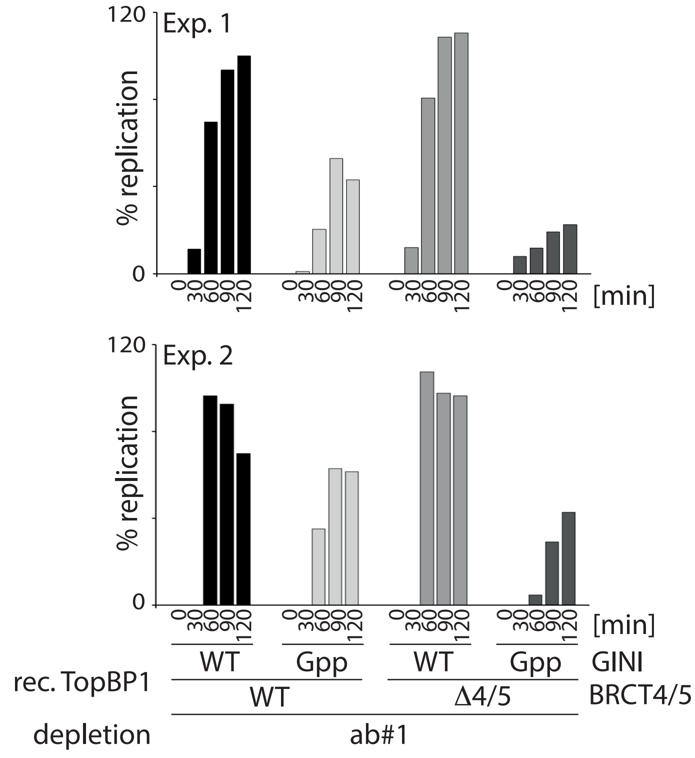

**b**

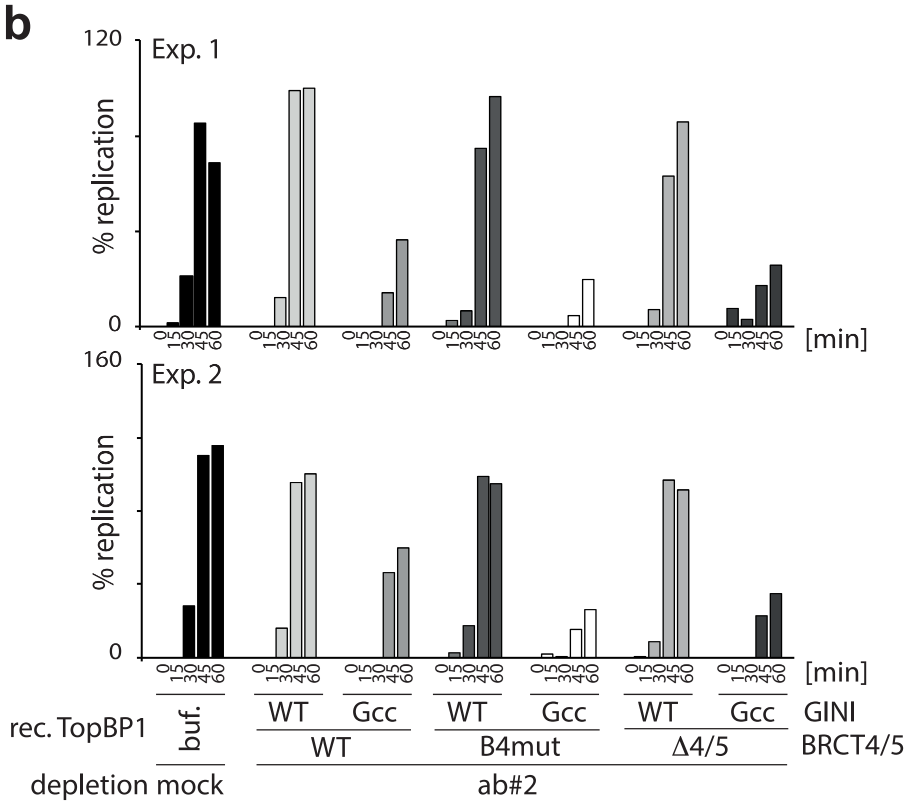

#### Supplemetary Figure 14

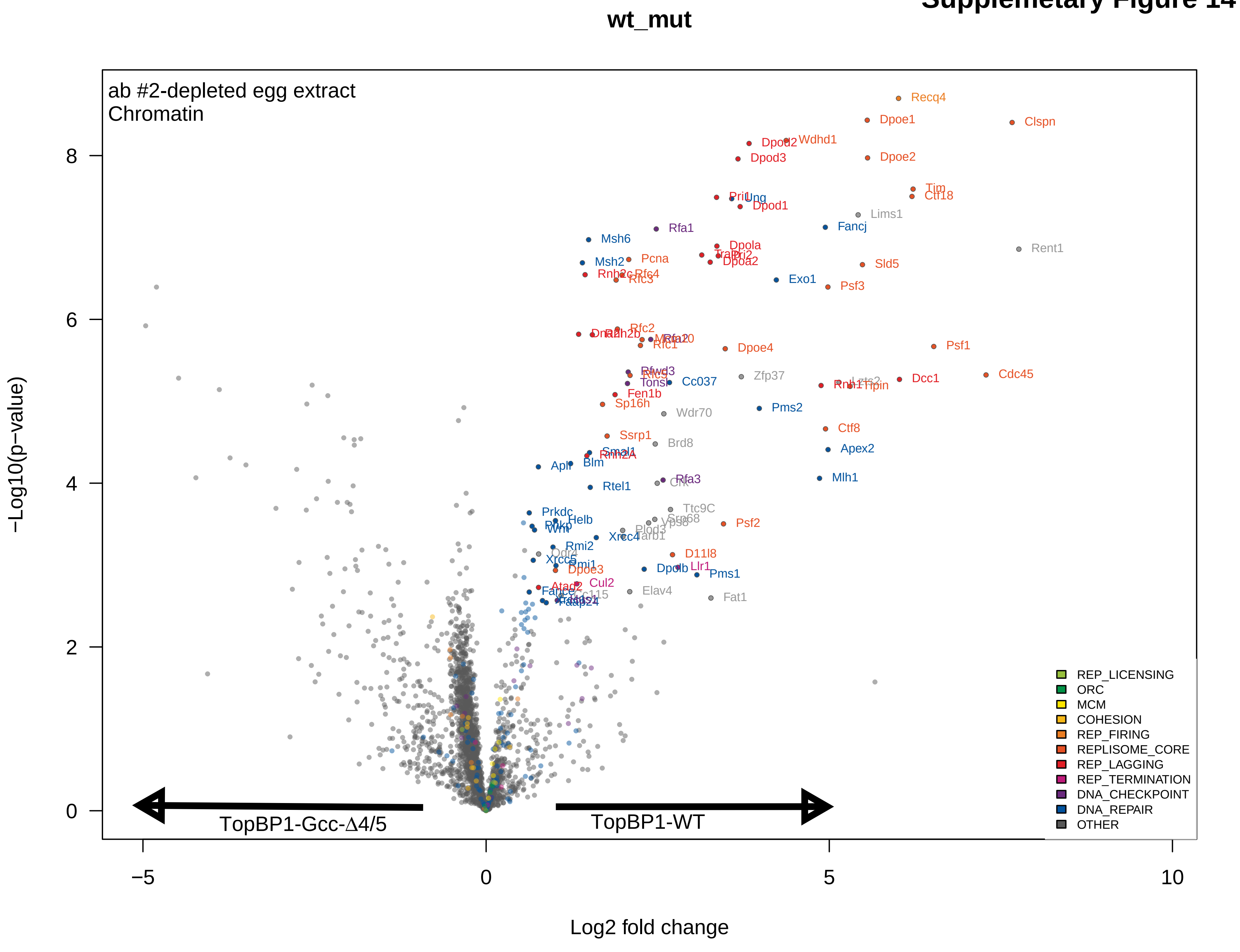

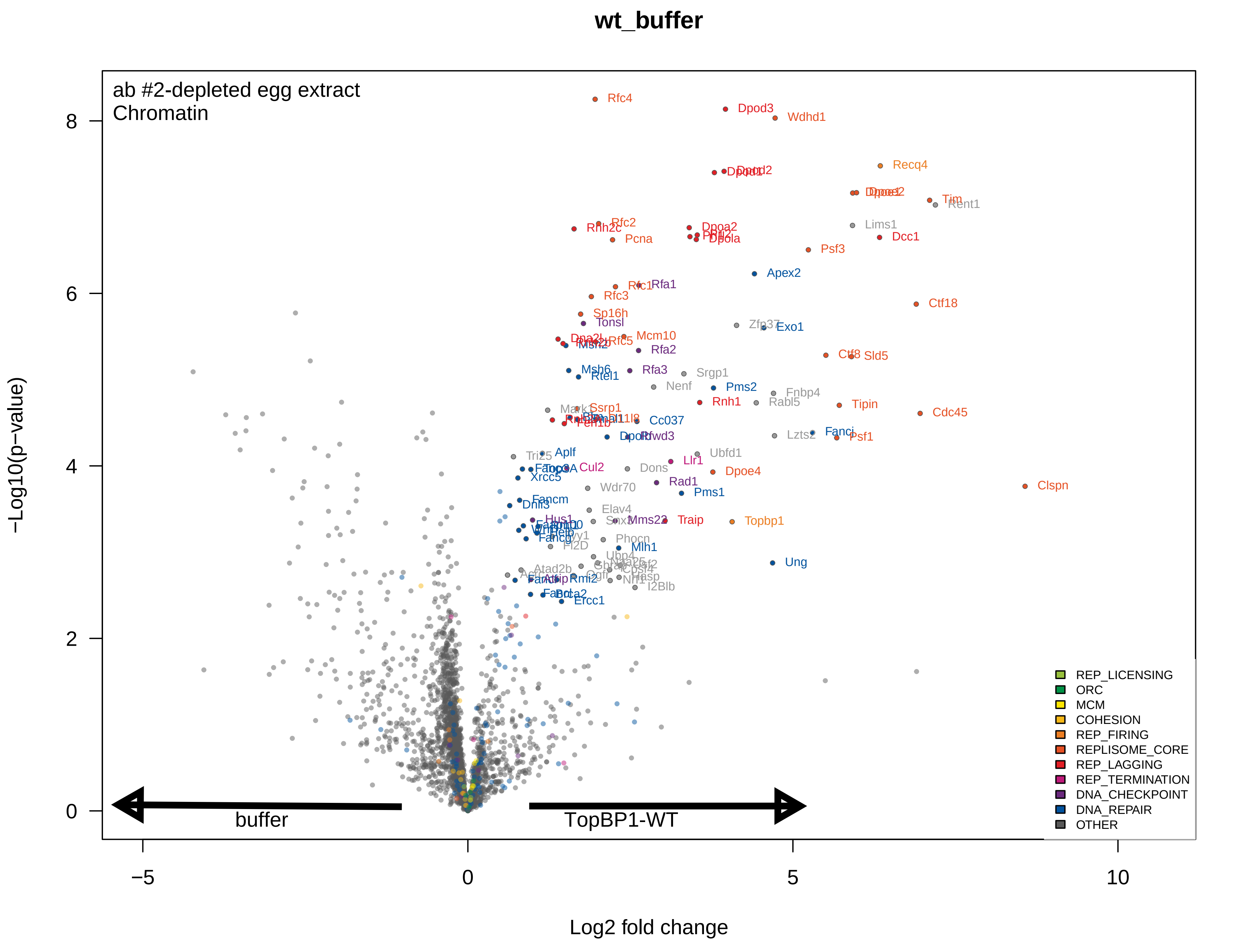
wt_buffer
ab #2-depleted egg extract
Chromatin
−Log10(p−value)
Log2 fold change
8
6
4
2
0
−5
0
5
10
buffer
TopBP1-WT
Rfc4
Dpod3
Wdhd1
Recq4
Tim
Lims1
Dcc1
Rfc2
Pcna
Psf3
Apex2
Rfa1
Rfc3
Ctf18
Sp16h
Tonsl
Exo1
Mcm10
Rfa2
Sld5
Msh6
Rtel1
Rfa3
Srgp1
Nenf
Pms2
Fnbp4
Rnh1
Rabl5
Tipin
Cdc45
Ssrp1
Cc037
Lzts2
Fanci
Psf1
Aplf
Ubfd1
Llr1
Cul2
Dons
Dpoe4
Xrcc5
Rad1
Wdr70
Pms1
Clspn
Fancm
Dnli3
Elav4
Mms22
Traip
Topbp1
Hus1
Fancg
Phocn
Mlh1
Ung
Atad2b
Rmi2
Ercc1
I2Blb
REP_LICENSING
ORC
MCM
COHESION
REP_FIRING
REPLISOME_CORE
REP_LAGGING
REP_TERMINATION
DNA_CHECKPOINT
DNA_REPAIR
OTHER

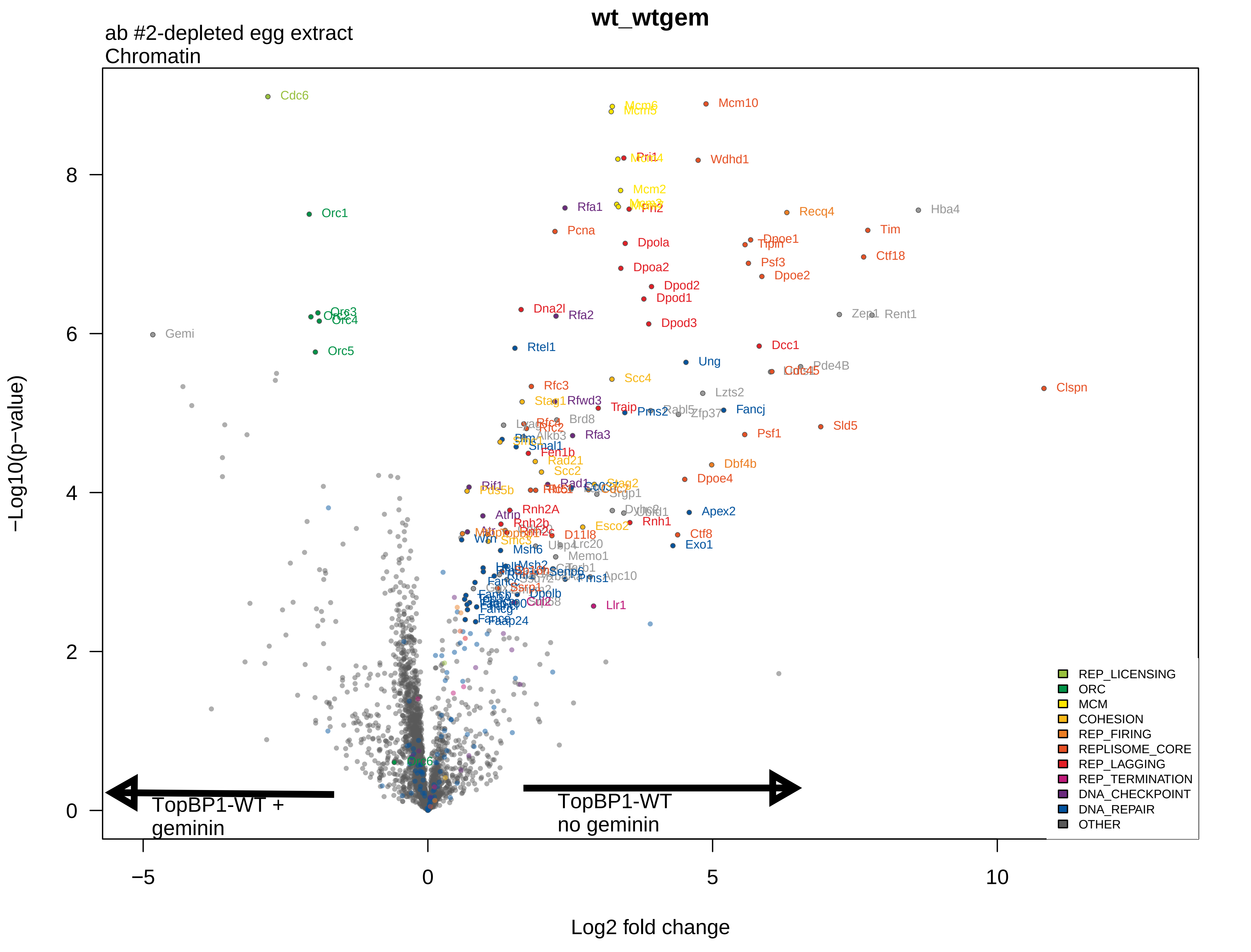

wt_wtgem
ab #2-depleted egg extract
Chromatin
−Log10(p−value)
Log2 fold change
−5
0
5
10
0
2
4
6
8
Cdc6
Mcm10
Wdhd1
Orc1
Rfa1
Recq4
Hba4
Pcna
Tim
Dpola
Ctf18
Psf3
Dpoa2
Dpoe2
Dpod2
Dpod1
Dna2l
Rfa2
Dpod3
Gemi
Orc5
Rtel1
Dcc1
Ung
Pde4B
Clspn
Rfc3
Scc4
Lzts2
Rfwd3
Traip
Fancj
Sld5
Psf1
Brd8
Rfa3
Fen1b
Rad21
Scc2
Dbf4b
Dpoe4
Rif1
Pds5b
Srgp1
Apex2
Rnh2A
Rnh1
Ctf8
Exo1
Esco2
Msh6
Memo1
Pms1
Apc10
Llr1
Faap24
Dpolb
Orc6
TopBP1-WT +
geminin
TopBP1-WT
no geminin
REP_LICENSING
ORC
MCM
COHESION
REP_FIRING
REPLISOME_CORE
REP_LAGGING
REP_TERMINATION
DNA_CHECKPOINT
DNA_REPAIR
OTHER

### Supplementary Figure 15

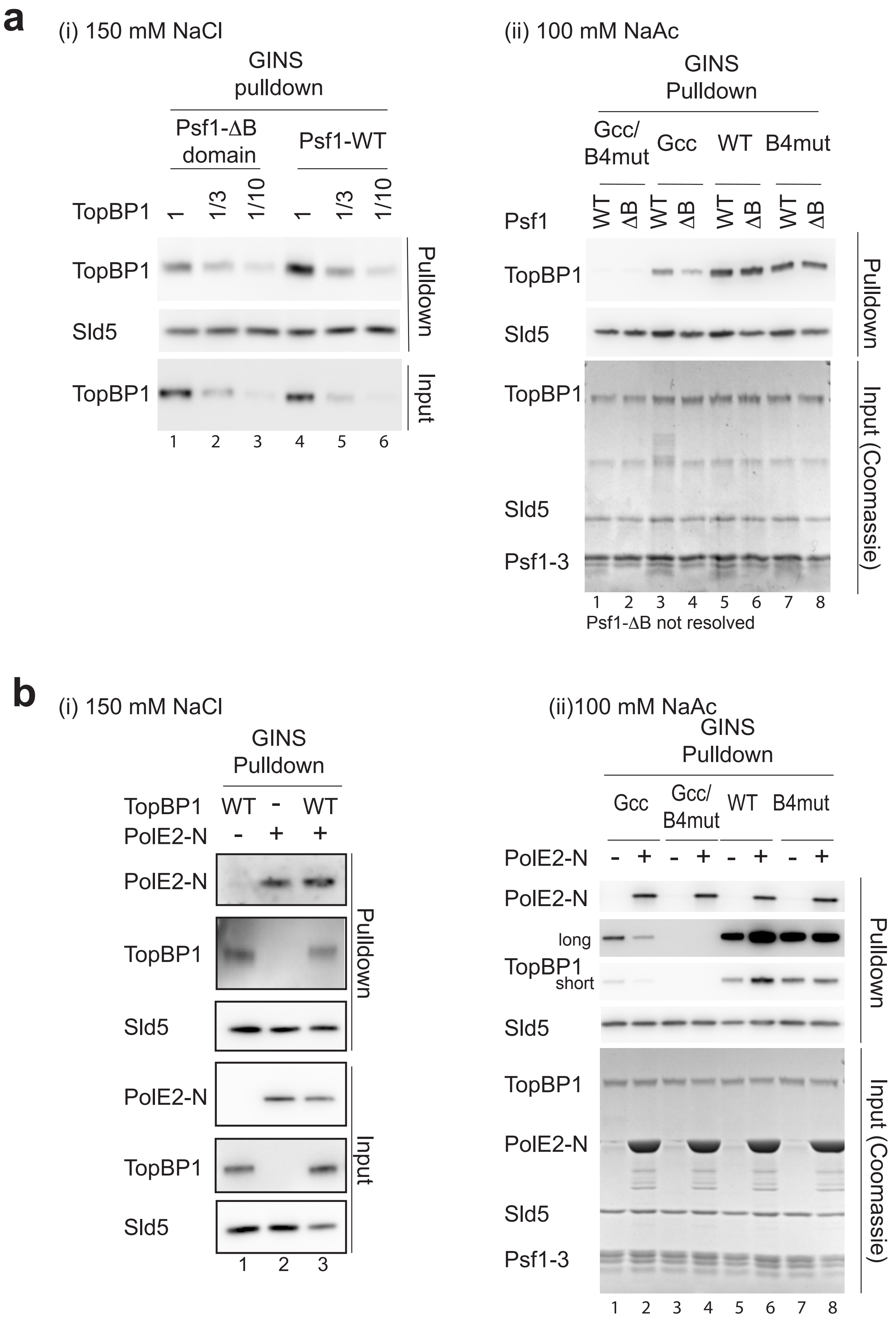

### Supplementary Figure 16

#### Figure S16

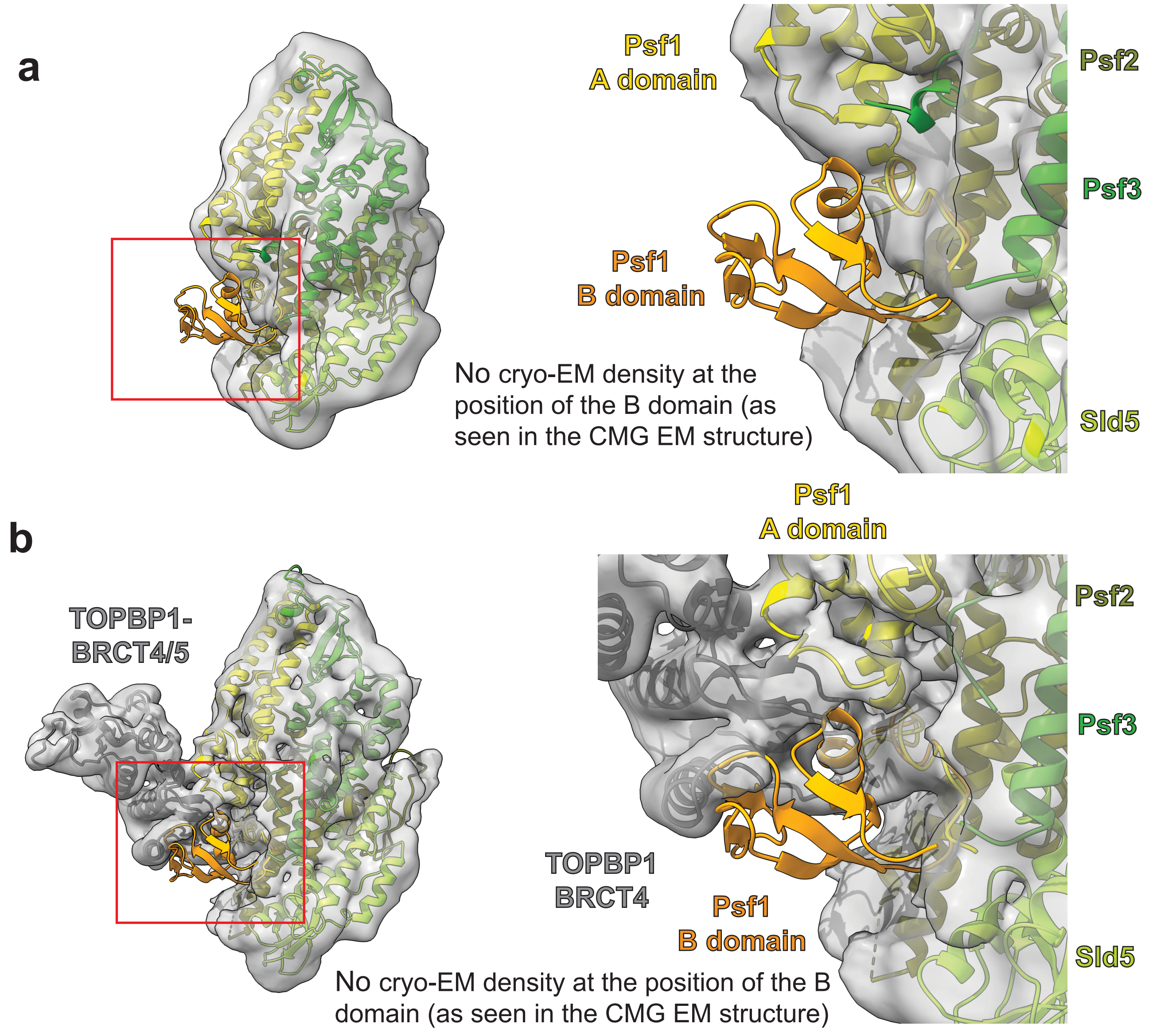

### Supplementary Figure 17

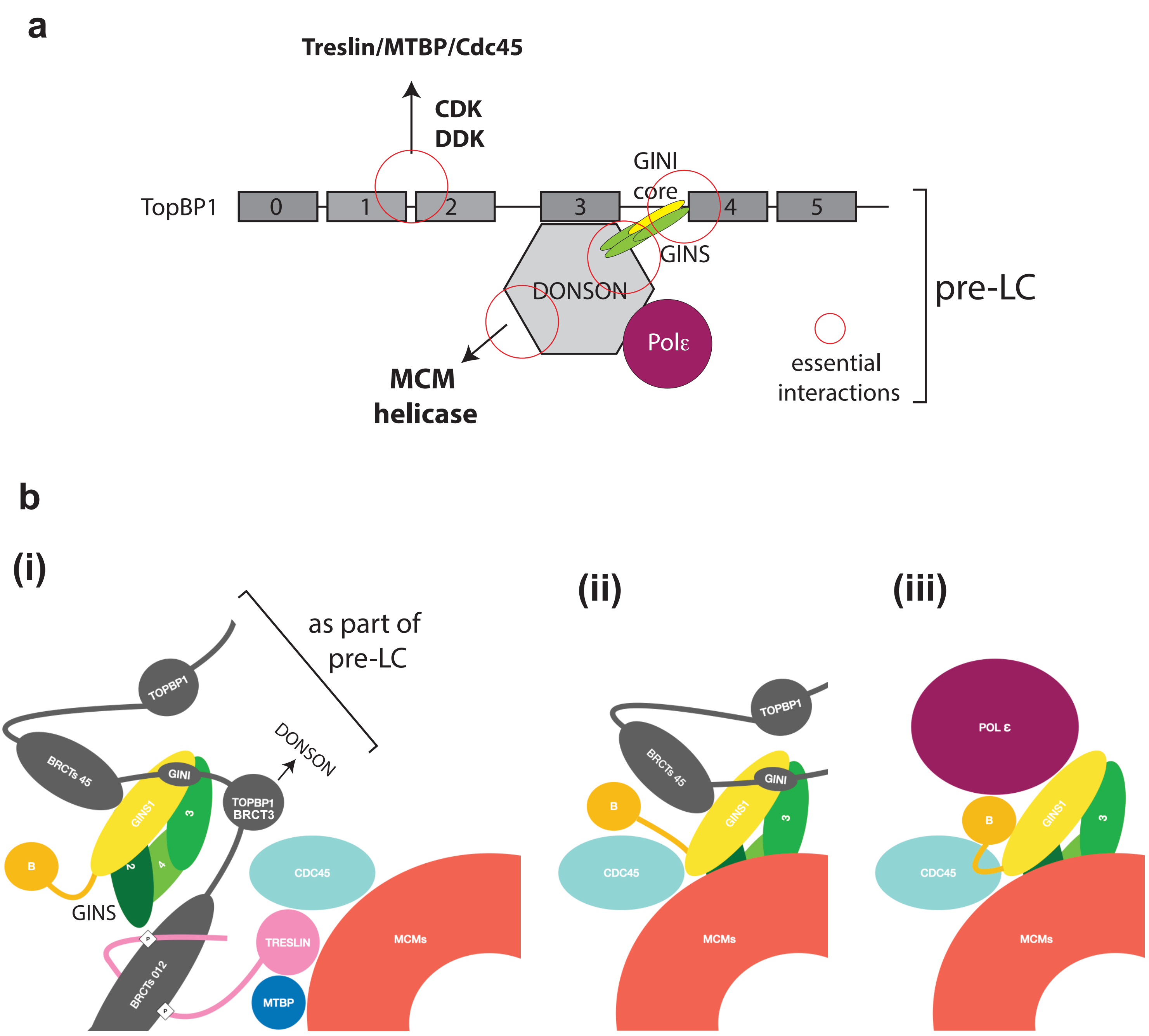

### Supplementary Table 1

**Cryo-EM data collection, refinement and validation statistics**

| | (EMD-16916)<br>(PDB 8OK2) |
|---|---|
| **Data collection and processing** | |
| Magnification | 120000 |
| Voltage (kV) | 300 |
| Electron exposure (e–/Å$^2$) | 39.69 |
| Defocus range (μm) | 1.2 – 2.2 |
| Pixel size (Å) | 0.74 |
| Symmetry imposed | C1 |
| Initial particle images (no.) | 13876268 |
| Final particle images (no.) | 154278 |
| Map resolution (Å) | 4.1 |
| FSC threshold | 0.143 |
| **Refinement** | |
| Initial model used (PDB code) | 29EX, 3UEN |
| Model resolution (Å) | 3.5 |
| FSC threshold | 0.143 |
| Model composition | |
| Non-hydrogen atoms | 7430 |
| Protein residues | 942 |
| Ligands | 0 |
| *B* factors (Å$^2$) | 78.33 |
| R.m.s. deviations | |
| Bond lengths (Å) | 0.002 |
| Bond angles (°) | 0.587 |
| Validation | |
| MolProbity score | 1.56 |
| Clashscore | 6.87 |
| Poor rotamers (%) | 0.49 |
| Ramachandran plot | |
| Favored (%) | 96.93 |
| Allowed (%) | 2.96 |
| Disallowed (%) | 0.11 |

#### Supplementary figure legends

**Supplementary Fig. 1: Schematic model of the origin firing reaction in yeast**
Details are described in the main text.

**Supplementary Fig. 2: Interaction of GINS and TopBP1 in cell lysates.**
a: Volcano plots showing label-free mass spectrometry quantification of proteins biotinylated by TopBP1 fused to APEX2. The top plot shows the results for stable isogenic Hek293-FlipIn cells while the middle and lower plots are based on results from transiently transfected Hek293T cells. Non-APEX2-biotinylated samples (-$H_2O_2$) were used as controls. Four biological replicates were analysed per experiment. Red dots and text indicate significantly enriched known TopBP1 interactors. Black dots and text indicate non-significantly enriched TopBP1 interactors.

b: Immunoblots showing immunoprecipitation of 6Myc-TEV2-TopBP1-BRCT0-5 from transiently transfected 293T cell lysates using anti-Myc antibodies. Non-transfected cell lysates were used for control IPs. Immunoprecipitated TopBP1 was detected using Ponceau (Ponc.) staining. A weak but specific co-IP of Sld5 was detectable.

c: Myc-immunoprecipitation as in b, except that 6Myc-TEV2-TopBP1-BRCT0-5-WT was compared with 6Myc-TEV2-TopBP1-ΔBRCT4/5. The specific co-IP of Sld5 with TopBP1 found in b could be confirmed. TopBP1-ΔBRCT4/5 bound GINS less efficiently compared to WT.

**Supplementary Fig. 3: Characterisation of the GINS-TopBP1 interaction.**
a: Coomassie-stained SDS gel of fractions from the size exclusion chromatography experiment testing complexation of TopBP1-BRCT0-5-strep and GINS tetramer shown in Fig. 1b. Where both GINS and TopBP1 were mixed there is a shift of the individual components to the earlier fractions indicating a stable complex was formed.

b: Recombinant TopBP1-ΔBRCT0-2 was incubated with recombinant GINS before size exclusion chromatography using a Superdex200 increase 10/300 column. The individual recombinant proteins were run separately for comparison. Elution profiles are shown with GINS coloured green, TopBP1 grey and the complex is coloured cyan. The clear shift upon complex formation suggests the N-terminal BRCT0-2 module of TopBP1 is dispensable for the interaction.

c: Recombinant GINS immobilised on Flag-beads and control beads (Flag beads coupled with Flag peptide) were used to pull down recombinant TopBP1-ΔBRCT3 (Fig 1a(ii)). SDS gel were silver stained.

d: The TopBP1 triple point mutant inactivating the phospho-binding pocket of BRCT5 (B5mut) that carries the mutations S654V, K661R and K704A (Fig. 1a(ii)) binds 53BP1 weakly in pulldowns from HEK293T cell lysates.

e: *In vitro* pulldown of TopBP1-WT and TopBP1-BRCT0-5-B5mut (Fig. 1a(ii)) with immobilised GINS (or Flag peptide for controls). The phospho-binding pocket of BRCT5 is inactive in B5mut (panel c). The B5mut mutant binds to GINS as well as TopBP1-BRCT0-5-WT, suggesting that the phospho-binding pocket of BRCT5 is dispensable for GINS binding.

**Supplementary Fig. 4: EM processing diagram for the initial cryo-EM dataset**

a: Data processing flowchart including an example micrograph, 2D class averages, and 3D classes with particle counts and resolution estimates at each step. Two volumes were produced from this dataset, one of the GINS complex alone (pink volumes and boxes) and one for the GINS-TopBP1 complex (blue volumes and boxes).

b: Orthogonal views of final 3D volume of the GINS-TopBP1 complex coloured by local resolution estimate.

c: Orientation distribution of particles included in final volume of the GINS-TopBP1 complex demonstrating preferred orientation but also that most views are represented by some particles.

d: FSC curves for final volume of the GINS-TopBP1 complex give an overall resolution of 4.6 Angstroms.

e: Orthogonal views of final 3D volume of the GINS complex coloured by local resolution estimate.

f: Orientation distribution of particles included in final volume of the GINS complex demonstrating preferred orientation but also that most views are represented by some particles.

g: FSC curves for final volume of the GINS complex give an overall resolution of 5.9 Angstroms.

**Supplementary Fig. 5: TopBP1-BRCT4 diverts from the canonical BRCT fold**

a) Superposition of BRCA1-BRCT1 (canonical BRCT; blue) and BRCT4 of TopBP1 (black). The canonical helix between beta strands three and four is missing in TopBP1-BRCT4.

b) Structural model of BRCT4 binding to Psf1. To show that the missing helix in BRCT4 helps create space for the interaction to take place, BRCT4 is shown here in the same relative orientation as in panel a. V590 and T606 are residues important for the interaction that were mutated in TopBP1-B4mut.

c) Multiple sequence alignment of the variant region in BRCT4 in a wide range of metazoa shows that the binding region for Psf1 constitutes a highly conserved patch. Residues mutated in TopBP1-B4mut show high similarity.

**Supplementary Fig. 6: Experiments to control quality of recombinant protein mutants.**

a: Recombinant TopBP1 point mutants of the GINI and BRCT4 regions bind 53BP1 phospho-dependently. TopBP1-BRCT0-5-strep-WT and indicated mutants (Figs. 1a and 3a summarize the TopBP1 mutants) were used to pull down endogenous 53BP1 from lysates of transiently transfected Hek293T cells. Lysates and pulldowns were analysed by immunoblotting. 53BP1 has two binding domains in TopBP1, BRCT0-2 and BRCT4/5 that both interact phosphorylation-dependently, as PPase-treatments showed (last two lanes). TopBP1-BRCT0-5 contains both binding sites, explaining its stronger binding to 53BP1 compared to the TopBP1-BRCT-ΔBRCT4/5 [42].

b: Psf1 I97R and Psf1 N43A/K51A form GINS tetramers normally. 100 nM of GINS-WT or the mutants were immobilized to the magnetic αFlag beads through 6xHis-3xFlag-Sld5 and then washed. Tetramer formation was detected by SDS PAGE and coomassie staining. GINS subunits were present in similar amounts, indicating normal GINS tetramer formation.

**Supplementary Fig. 7: Alignment of TopBP1 proteins from different vertebrate species using T-COFFEE and relative position to the TopBP1 BRCT domains.**
Xenopus, *Xenopus laevis* (Q7ZZY3), Cow, *Bos taurus* (A0A3Q1LWE4), Rat, *Rattus norvegicus* (A0A8I6GFZ6). Coloured labelling shows the amino acids mutated in the TopBP1 region I (RI), region II (RII), region III (RIII), and GINI core centre (Gcc) mutants. ., : and * indicate low conservation, high conservation and identical amino acids, respectively.

**Supplementary Fig. 8: Probability plots for AlphaFold2-Multimer prediction of Fig. 4a.**
PAE and pLDDT plots are shown for 5 models produced for TopBP1-BRCT3-5 complexed to the GINS tetramer predicted using AlphaFold2-Multimer. In all five models both interaction surfaces of TopBP1 can be seen to have low PAE scores with regards to the GINS subunits. Model 1 was used to produce the structural images in Fig. 4a.

**Supplementary Fig. 9: Molecular docking simulations** of the GINI centre core peptide (487-DEDLLSQY-494) on the GINS tetramer using Cluspro (a), MDockPep (b) and HPEPDOCK (c). Docking refinement with FlexPepDock (d).

**Supplementary Fig. 10: Crosslinking mass spectrometry with GINS-TopBP1.**

A: Schematic representing $BS^3$-mediated cross-links between TopBP1-BRCT0-5-Strep and GINS: (i), crosslinks involving the TopBP1-GINI domain, (ii), all inter-protein crosslinks. Recombinant TopBP1-BRCT0-5-strep (amino acids 1-766) and GINS crosslinked with 600µM or 2500µM of $BS^3$ crosslinker were analysed by mass spectrometry, both in solution (left) and after SDS-PAGE and in-gel digestion (right). ProXL (Protein Cross-Linking Database) was used to analyse and visualize the data. Crosslinks in (i) are coloured according to the number of peptide spectrum matches showing this link.

b: AlphaFold2 structure prediction of the TopBP1-Psf1-Psf3 complex with violet highlighted amino acids involved in crosslinking in panel a.

c: GINS crystal structure superposed onto AlphaFold2 TopBP1-Psf1-Psf2 complex with Sld5 amino acids involved in crosslinking highlighted in violet.

GINI region:
GINI region crosslinks mostly clustered, involving Psf1 lysines 63, and Psf3 Lys80. Fewer crosslinks involved Psf1 Lys61 situated in the same region but facing the other direction. The distance between Lys482 in the GINI helix and Thr61 of Psf1 is 27,25 Å. Between TopBP1-Ser480 and Psf3-Lys74, it is 26,16 Å. The largest detected distance of 50 Å involving TopBP1 lysines 466 and 468 is probably explained by the flexibility of this unstructured part, which is supported by crosslinking with Psf3 lysine 65 on the other face of this structural element.

Other TopBP1 and GINS regions:
The GINI region also crosslinks with Sld5 residues 112, 113, 142, 164 and 208 situated in the vicinity of the predicted GINI interaction site in Psf1/3 (compare panel c).
The GINI region appears a crosslinking focus in TopBP1. Further crosslinks involve TopBP1-BRCT0-2, mainly with Sld5 (residues 102, 112, 113, 142, 164, 208 and 209) and Psf1 (residues 51, 61, 63, 154 and 171).
Surprisingly, BRCT4-GINS crosslinking is not apparent, with only one crosslink detected (BRCT4 residue 576 and Psf3 residue 210). Perhaps, the BRCT4 binding region of Psf1 is crosslinked in a conformation that prevents BRCT4 binding. The detected crosslinking between the C-terminal helix of Psf3 and the Psf1 B domain (reminiscent of confirmation in CMG) might interfere with TopBP1 BRCT4-GINS complexation.

**Supplementary Fig. 11: EM processing diagram for the second dataset, using Psf1-ΔB (B domain deleted**

a: Data processing flowchart including an example micrograph, 2D class averages, and 3D classes with particle counts and resolution estimates at each step.

b: Orthogonal views of final 3D volume coloured by local resolution estimate.

c: Orientation distribution of particles included in final volume demonstrating preferred orientation but also that most views are represented by some particles.

d: FSC curves for final volume give an overall resolution of 4.1 Angstroms.

**Supplementary Fig. 12: Characterisation of TopBP1 mutants**
a: Pulldowns of the indicated TopBP1 mutants with bead-immobilised GINS or Flag peptide-coupled control beads at 50 mM, 100 mM and 200 mM sodium. Whereas pulldowns in 150 mM NaCl did not show detectable binding of the BRCT4 and GINI regions (Fig. 2c, 3b) 50 and 100 mM sodium acetate revealed appreciable binding at reduced levels compared to TopBP1-WT. TopBP1 double mutants did not pull down with GINS.

b: Immuno-depletion and add-back experiment as described in Figures 5e/f and 6. Western blot with the indicated antibodies upon re-isolating sperm chromatin from egg extract 120 minutes after starting the replication reaction. TopBP1-Gcc mutant shows slightly stronger signals than the double mutant for Cdc45, but clearly weaker than WT and Δ4/5.

**Supplementary Fig. 13: Replication analyses using *Xenopus* egg extracts**
a: Replication analysis of experiment as described in Fig. 5e/f, but using anti-TopBP1 antibody #one. Two independent experiments (each n=1) are shown.

b: Replication analysis using *Xenopus* egg extracts upon replacing endogenous TopBP1 with indicated recombinant TopBP1 versions to test the consequences of the TopBP1-B4mut triple point mutations that were identified by our structural studies (Fig. 2b,c). TopBP1 antibody #2 was used. TopBP1-B4mut and Δ4/5 behave similarly. The mutations in B4mut do not affect grossly the replication-inducing capacity compared to TopBP1-BRCT0-5-WT. However, the mutations in B4mut aggravate the replication defect by the GINI site mutant Gcc. This aggravation effect is consistent with the model that the residues in BRCT4 that are required for normal GINS binding are also required for the normal replication-inducing activity of TopBP1. The time points shown in this experiment are different from the others in this study, because the egg extract used here replicated quicker. Two independent experiments (each n=1) are shown.

**Supplementary Fig. 14: Volcano plots of mass spectrometry-based chromatin analysis**
Data of experiments described in Figure 6a/b is shown. Here all differentially detected proteins are labelled (see Materials and Methods and supplementary table 2 for details).

**Supplementary Fig. 15: Interaction studies support that the Psf1 binding sites for PolE2-N terminal domain and TopBP1-BRCT4 partially overlap.**
a: The Psf1-B domain is required for full affinity binding of TopBP1 in vitro. a) Pulldown experiments comparing the binding of bead-immobilised GINS-WT and mutant GINS containing Psf1 lacking the B domain (Psf1-ΔB) to TopBP1-0-5-WT and the indicated mutants. An experiment with TopBP1-WT in stringent conditions (150 mM NaCl) under (i) shows

dispensability of the Psf1-B domain for TopBP1-WT interaction. (ii) shows an experiment in lower ionic strength (100 mM NaAc) testing TopBP1-WT, Gcc and B4mut, which demonstrates that the B domain deletion affects binding to BRCT4 but not to WT and Gcc.

b) Excess of PolE2-N terminal domain competes TopBP1-Gcc, but not TopBP1-0-5-WT and B4mut off of bead-immobilised GINS. Bead-immobilised GINS was pre-incubated with ten times molar excess of MBP-tagged PolE2-N terminal domain (amino acids 1-75) over TopBP1. After ten minutes TopBP1-WT or mutants were added for the subsequent pulldown. PolE2-N partially competes TopBP1-Gcc but not the other TopBP1 versions off of GINS, showing partial competition between PolE2-N and the BRCT4

**Supplementary Fig. 16: The Psf1-B domain is not visible in our cryo-EM data of the GINS tetramer and the GINS-TopBP1 complex.**

a: GINS alone map volume with docked crystal structure of GINS (PDB:2E9X) shown alongside the B domain of Psf1 taken from a superposed model of the full GINS complex from a cryo-EM reconstruction of the full CMG structure (PDB:6XTX) demonstrating no density for the B domain of Psf1 can be seen when the GINS complex is free suggesting a flexible conformation.

b: GINS-TopBP1 volume shown with TopBP1-GINS model from this paper, alongside the B-domain of Psf1 as for panel a, showing in our dataset no density for the B domain of Psf1 can be seen when GINS is bound to TopBP1. Taken together these two volumes suggest the B domain of Psf1 is flexible when not bound to the MCM ring and CDC45.

**Supplementary Fig. 17: Models of TopBP1-mediated loading of GINS onto MCM-DHs.**

a: TopBP1 functions as a regulated scaffold within the pre-LC context. TopBP1's numerous interaction domains are key to assemble pre-LC. The CDK and DDK controlled interaction with Treslin couples CMG formation to S phase. Here we demonstrate and characterise the interaction with GINS, and show that, collectively, the two GINS binding sites in TopBP1 are essential for origin firing. A second scaffold, DONSON, is an integral part of vertebrate pre-LC, whereas RecQL4 as a sequence homologue of yeast pre-LC component Sld2 is not part of vertebrate pre-LC. An interesting situation is present in pre-LCs of the metazoan *C. elegans* that possesses a yeast-like Sld2 in addition to DONSON.

b: Hypothetical model of TopBP1's role in the CMG formation process. TopBP1 is recruited through a phosphorylation dependent interaction between the N-terminal BRCT0-2 module and Treslin (i). This allows delivery of the GINS tetramer to the MCM-CDC45 complex (ii). The Psf1-B domain orders, packing against CDC45, simultaneously locking the MCM ring and weakening the interface between TopBP1 BRCT4 and GINS complex (iii). TopBP1 dissociates, exchanging with Polε to form the CMGE complex.

#### Supplementary table legends

**Supplementary Table 1:**
**Cryo-EM data collection, refinement and validation statistics**
